# High efficiency transfection of thymic epithelial cell lines and primary thymic epithelial cells by Nucleofection

**DOI:** 10.1101/030221

**Authors:** Richard T. O’Neil, Qiaozhi Wei, Brian G. Condie

## Abstract

Thymic epithelial cells (TECs) are required for the development and differentiation of T cells and are sufficient for the positive and negative selection of developing T cells. Although TECs play a critical role in T cell biology, simple, efficient and readily scalable methods for the transfection of TEC lines and primary TECs have not been described. We tested the efficiency of Nucleofection for the transfection of 4 different mouse thymic epithelial cell lines that had been derived from cortical or medullary epithelium. We also tested primary mouse thymic epithelial cells isolated from fetal and postnatal stages. We found that Nucleofection was highly efficient for the transfection of thymic epithelial cells, with transfection efficiencies of 30-70% for the cell lines and 15-35% for primary TECs with low amounts of cell death. Efficient transfection by Nucleofection can be performed with established cortical and medullary thymic epithelial cell lines as well as primary TECs isolated from E15.5 day fetal thymus or postnatal day 3 or 30 thymus tissue. The high efficiency of Nucleofection for TEC transfection will enable the use of TEC lines in high throughput transfection studies and simplifies the transfection of primary TECs for *in vitro* or *in vivo* analysis.

## Background

The non-hematopoietic epithelial cells of the thymus form a complex microenvironment that supports the normal development and differentiation of T cells. Developmentally, thymic epithelial cells (TECs) are derived exclusively from the pharyngeal endoderm [1]. During thymus organogenesis TEC progenitors differentiate into medullary or cortical cell types and these two lineages form histologically and functionally distinct compartments within the thymus [2]. Interactions between the non-hematopoietic TECs and differentiating T-cells are required for normal T cell differentiation, maturation and selection [3]. Due to their essential role in T cell development, understanding the process of thymic epithelial cell specification, differentiation and maintenance will be required to understand T cell biology. TECs are also of clinical interest due to the progressive loss of thymus function during normal human aging. Slowing or reversing the loss of TECs may prolong robust T cell function in aging individuals. In addition, a greater understanding of thymic epithelial cell specification and differentiation during development may pave the way for the generation of TEC stem cells that can be transplanted to restore or supplement endogenous thymic function in ageing individuals or those with congenital thymus dysgenesis or agenesis [4-6].

Given the importance of thymic epithelial cells in basic and clinical immunology several approaches have been developed for their analysis *in vivo* and in cell culture. Genetic studies in mice and humans have identified a number of genes required for TEC formation, differentiation and function [7-11]. In addition, cell labeling, tissue grafting, cell culture, and organ culture approaches have been developed for studies of embryonic TEC lineage specification, differentiation or TEC-thymocyte interactions [12-15]. Primary thymic epithelial cell cultures can be readily established from fetal or adult thymus and many thymic epithelial cell lines are available. Unfortunately, it is quite difficult to transfect primary TECs or established TEC lines using simple approaches such as lipofection. This difficulty has been noted previously [16] and many studies of TEC gene promoters or TEC transcription factors have been performed in HELA or other non-TEC cell lines [16-21]. Although methods for TEC transfection via retroviral vectors [22] have been developed, these methods are cumbersome since they require construction of viral vectors and the production of high titer stocks for each construct. This makes viral approaches problematic in cases where a large number of independent constructs must be tested such as in promoter analysis or in studies of protein functional domains.

To remove this current limitation to the experimental analysis of TEC biology, we tested whether nucleofection can efficiently transfect plasmid constructs into 4 independent TEC lines as well as primary TECs isolated from three different developmental stages. Nucleofection is a relatively new highly optimized electroporation method that enables high efficiency electroporation of a wide range of cell types including primary cells and postmitotic cells [23]. Nucleofection has been shown to efficiently transfect a wide range of cell types including stem cells, progenitors and/or fully differentiated cells isolated from the nervous system, the immune system, smooth muscle, skeletal muscle, adipose tissue, skin, connective tissue, cartilage, bone marrow stroma, vascular endothelia as well as mouse and human embryonic stem cells [23-45]. Many of the cell types that can be efficiently transfected by nucleofection are extremely difficult or impossible to transfect using other non-viral methods.

Given Nucleofection’s track record, we tested whether it would be an efficient method for the routine transfection of thymic epithelial cells. We found that nucleofection does result in high efficiency transfection of TEC lines as well as primary TECs. We also found that reproducible high efficiency transfection was possible while preserving cell viability. Our studies indicate that nucleofection greatly increases the feasibility of studies that require the transfection of multiple independent constructs into thymic epithelial cells. This simple and high efficiency transfection method will facilitate the analysis of gene regulatory regions of TEC genes as well as the analysis of the functional domains of proteins that are necessary for TEC development and function.

## Results and Discussion

### Thymic epithelial cell lines can be transfected at high efficiency via Nucleofection

To test whether Nucleofection could be used to efficiently transfect a range of different thymic epithelial cell lines we examined the Nucleofection efficiency of four different TEC lines (TE-71, Z210R.1, 100-4, 41.2). The TE-71 and Z210R.1 TEC lines were derived from normal adult Balb/c mice and were typed as medullary epithelium [46]. The 100-4 and 41.2 cell lines were derived from C3HXC57B1/6 mice and were derived from cortical epithelium [47, 48].

Our initial goal was to test several different Nucleofector programs to define an optimal program for each of the four cell lines tested. To do this we used a reagent kit that had been recommended by Amaxa support staff for the optimization of conditions for epithelial cell lines (kit V). For each cell line we measured the transfection efficiency and cell survival with the same set of 6 different Nucleofector programs. Using this approach, we identified conditions for the high efficiency transfection of all four TEC lines with low cell mortality. For each Nucleofector program we found that the maximum transfection efficiency differed between the cell lines. However, we found that for all cell lines, program T-30 resulted in the most efficient transfection with the lowest amount of cell death (Figure 1 A, B).

**Figure 1.**
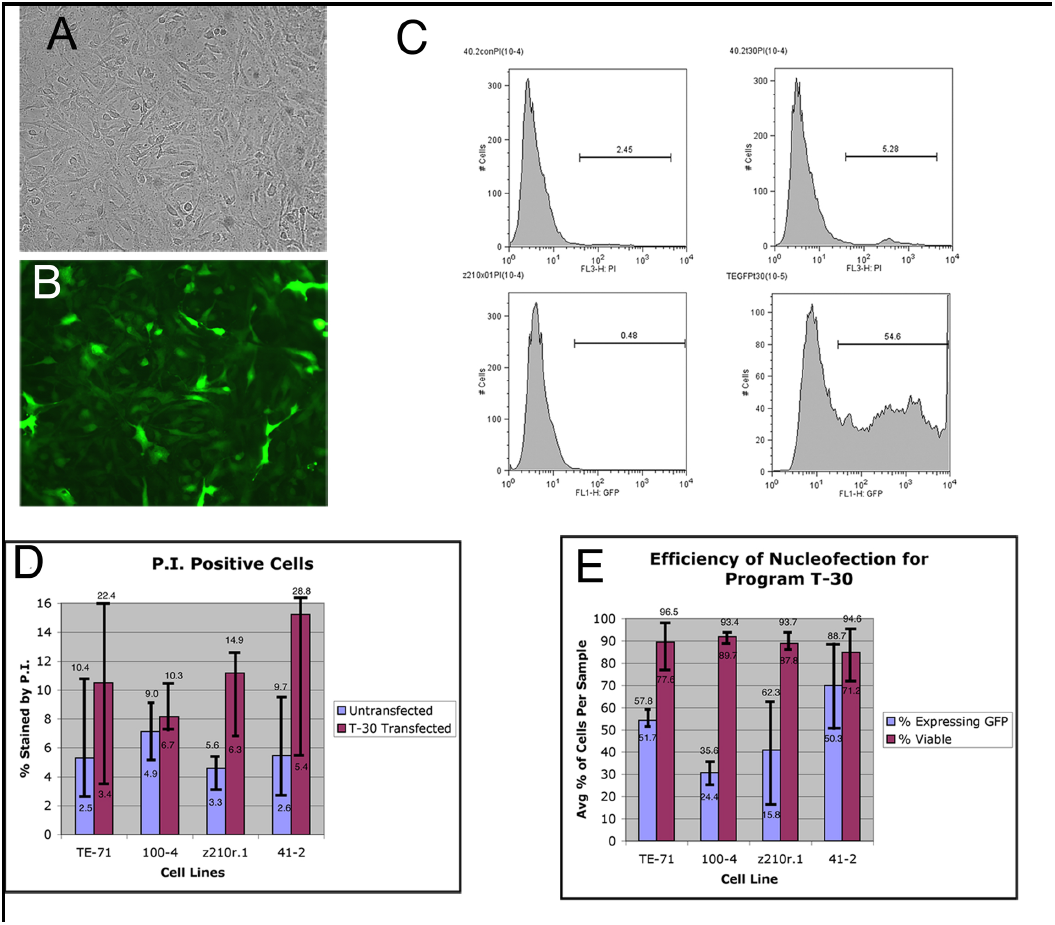
Efficient transfection of four different thymic epithelial cell lines by Nucleofction. (A, B) GFP expression (B) 24 hours after Nucleofection (program T-30) of the pmaxGFP plasmid into the thymic epithelial cell line 41–2 which was derived from cortical thymic epithelium; (A) brightfield phase-contrast image of the same field shown in B. (C) Results of flow cytometry performed 24 hours after Nucleofection. The data shown are representative results for a single experiment with the 41–2 TEC line and Nucleofection program T-30. The top row of plots show propidium iodide staining profiles for the untransfected (left panel) and transfected (right panel) cells. The bottom row shows the GFP staining profiles for untransfected (left panel) and pmaxGFP transfected (right panel) 41–2 cells. (D, E) Overall cell death as measured by propidium iodide staining (D) and the percentage of the cells expressing GFP the four TEC lines tested (E). Bars indicate the range of the average values obtained by flow cytometry.

Once we had identified T-30 as an optimal Nucleofection program for the TEC lines we then performed additional experiments to determine the reproducibility of TEC line Nucleofection. We measured the transfection efficiency as the percentage of GFP positive cells that had survived for 24 hours after Nucleofection by flow cytometry (Figure 1C). We found that our Nucleofection conditions resulted in mean transfection efficiencies of 30% to 70% for the four TEC lines. (Figure 1E). We also found that the reproducibility of Nucleofection efficiencies varied between the 4 cell lines with the range of the efficiency being smaller for the TE-71 and 100-4 cell lines than for the Z210r.1 and 41-2 cell lines (Figure 1E). Cell survival in all cases was excellent, with greater than 85% of the cells surviving for 24 hours after Nucleofection of each cell line (Fig. 1D). We also tested whether the transfection efficiency would vary with cell density. We found that Nucleofection of fewer than 1.5 × 10^6^ cells from each TEC line resulted in greatly reduced cell survival immediately after Nucleofection and that cell survival greatly improved as the sample size reached 4 × 10^6^ cells with no loss of efficiency (data not shown). Excellent viability and proliferation were observed for each of the four cell lines for up to 7 days after the procedure.

Primary TEC cultures are also used for studies of thymic development and function [49, 50]. Primary TEC cultures can be established from fetal as well as adult thymus. Unfortunately, TECs are a tiny minority of the cells in the thymus making it difficult to isolate large numbers of cells for primary culture. Therefore, high efficiency transfection methods that produce minimal cell mortality are an absolute necessity for experiments with primary thymic epithelial cells. We started our tests of primary TEC Nucleofection with cells isolated from E15.5 day fetal thymuses. At this stage there is a relatively large number of proliferating TECs while T-cell number is relatively low [51]. In an initial experiment to identify optimal Nucleofection conditions for primary fetal TECs, we tested several Nucleofector programs. The programs T-13, T-20 and U-17 resulted in high levels of cell death with less than 10% of the cells surviving the procedure. However, we found that there was a trade-off between cell death and transfection efficiency, with programs T-13 and U-17 resulting in about 50% of the surviving cells being GFP+. In contrast, transfection with program S-05 resulted in greater than 30% cell survival the with about 30% of the surviving cells being GFP+ (Figure 2, A-C). To quantify the absolute efficiency of each program, the total number of GFP expressing cells was taken as a ratio of the total number of viable cells in an untransfected control (Figure 2 C). Using this analysis it was determined that, of the programs tested, program S-05 was the most efficient program for Nucleofection of E15.5 mouse fetal primary TECs using the Nucleofector solution V.

**Figure 2.**
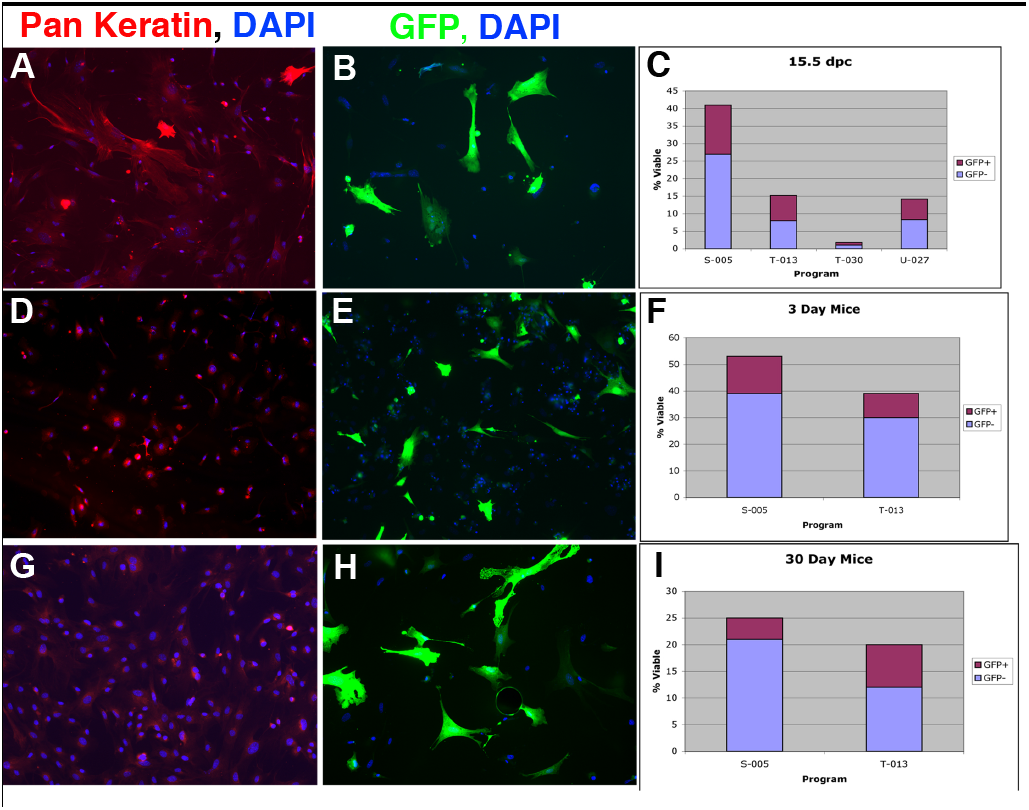
Efficient transfection of primary thymic epithelial cells by nucleofection. Thymic epithelial cell cultures were isolated from E15.5 day embryos, 3 day postnatal pups, and 30 day old mice. (A, D, G) Detection of cytokeratin by a pan-cytokeratin antibody reveals that most of the cells in the primary cultures are TECs. Primary TECs from E15.5 (A), postnatal day 3 (D) and postnatal day 30 (G) animals are shown. (B, E, H) Expression of the pmaxGFP plasmid in transfected primary TECs from E15.5 (B), postnatal day 3 (E) and postnatal day 30 (H) animals are shown. (C, F, I) Results from representative Nucleofections of primary TECs. Each bar on the bar graph shows the proportion of cells that survived Nucleofection as compared to untransfected control cultures. Each bar is divided into the proportion of cells expressing GFP (red segment of each bars) as well as the proportion that did not express GFP (blue segment of each bar. In the case of primary TECs derived from E15.5 fetal thymi the transfection results are shown for Nucleofection programs S-005, T-013, T-030 and U-027 (C).

Upon determination that program S-05 gave the highest transfection efficiency with good cell viability, we performed additional transfections to determine the reproducibility of this protocol with E15.5 day primary TECs. It was found that consistent levels of viability and GFP expression could be achieved with program S-05 with values similar to that found in the optimization experiments (data not shown). Cytokeratin immunostaining confirmed that nearly all of the cells in the cultures expressed cytokeratin, indicating that they are TECs (Figure 2A).

We also tested the efficiency of Nucleofection for the transfection of primary TECs isolated from postnatal thymuses. We tested the Nucleofection programs T-13 and S-05 on TEC cultures prepared from 3 day old (P3) and 30 day old (P30) postnatal Swiss Webster mice (Figure 2). We found that the P3 TECs were efficiently transfected, with about 25% of the viable cells expressing GFP (figure 2 D-F). However, we found that the nucleofection conditions had to be modified for the older primary TECs isolated from 30 day old thymuses. With the earlier stages we had used Nucleofection solution V, but for 30 day old postnatal primary cells use of this solution resulted in extremely low transfection efficiencies of around 1% of the viable cells (data not shown). To improve the transfection efficiency for the TECs from 30 day old mice we tested the primary epithelial cell Nucleofection solution with this cell type. We found that using program T-13 we were able to get high efficiency transfection with about 40% of the viable cells expressing GFP but with only 20% of the starting cell population remaining viable (figure 2 G-I). Although nucleofection of these later stage primary TECs results in greater cell death, the efficiency is quite high. Again, staining with pan-cytokeratin confirmed that the cells are TECs (figure 2G).

Our results show that Nucleofection enables the high efficiency transfection of primary TEC as well as cell lines derived from thymic epthelial cells. This cell type is difficult to transfect by methods typically used for routine transfection such as lipofection. A previous report described an efficient method for retrovirus-mediated transfection of primary TECs [22]. This method results in stable transfection of about 30% of the E15.5 TECs used in the study [22]. Retroviral transfection is an excellent choice for the generation of stable transfected cell lines but it is cumbersome for routine transient transfection studies. Retroviral vector production for TEC transfection requires the transfection of the viral packaging cell line, enrichment of the transfected cells and centrifugation to concentrate the viral particles [22]. Since most studies that use transient transfection utilize many expression and/or reporter constructs, retroviral transfection of TECs would be laborious and expensive.

In contrast, the high efficiency achieved by Nucleofection of TECs makes it feasible to perform any type of analysis that requires repeated transfections of many plasmid constructs. In comparison with lipofection we find that nucleofection results in greater cell viability and much greater transfection efficiency. For example, in our lab lipofectamine-mediated transfection of TECs results in a transfection efficiency of 3-5% using the same TEC lines that we used in this study (data not shown). Our finding that Nucleofection can transfect thymic epithelial cells with high efficiency and relatively low cell mortality adds another method to the rapidly developing molecular genetic toolkit for studies of thymic epithelial cells. Existing genetic tools for the manipulation of TEC biology include retroviral transfection methods as well as mouse strains that express GFP, Cre recombinase and cyclin D1 in all or some TEC populations and the identification of loci that can be used to drive gene expression in all or a defined subset of the TECs *in vivo* [22, 52-57]. By applying Nucleofection to the study of TEC biology we have enabled additional avenues for the molecular analysis of the development and function of this crucial cell type in cell culture.

## Conclusions

Thymic epithelial cells can be readily transfected at high efficiency by Nucleofection. Efficient transfection can be performed with established cortical and medullary thymic epithelial cell lines as well as primary TECs isolated from E15.5 day fetal thymus or postnatal day 3 or 30 thymus tissue. Transfection efficiencies as high as 70% of the surviving cell population can be achieved reproducibly. Nucleofection will enable high throughput transfection-based studies of TEC gene regulation, cell biology and differentiation.

## Methods

### Culture of TEC lines and the preparation and culture of primary TECs

The four TEC lines used in this work (100-4, Z210R.1, TE-71, 41.2) were obtained from Dr. Andrew Farr, Dept. of Immunology, University of Washington School of Medicine. All TEC lines were cultured on T-75 tissue culture flasks (Nunc) in RPMI 1640 (Invitrogen) medium with 10% fetal bovine serum (Invitrogen), 2 mM L-Glutamine (Invitrogen), 50 units/ml Penicillin and 50 μg/ml Streptomycin (Invitrogen). Each cell line was passaged into 2 T-75 flasks at 4 X10^6^ cells per flask and incubated at 37°C in 5 % CO2 for 48 hours prior to Nucleofection. After 48 hours of incubation, cells were 80-90% confluent and each flask contained 8 × 10^6^ to 1.4 × 10^7^ cells. Cells were trypsinized with 0.25% trypsin/EDTA (Invitrogen) and resuspended in aliquots of 2 × 10^6^ cells per 2 ml of media for nucleofection.

To generate fetal primary TEC cultures, thymi were removed from E15.5 C57Bl/6 embryos and placed directly in ice-cold TEC-10 media containing RPMI 1640 (Invitrogen) medium with 10% fetal bovine serum (Invitrogen), 2 mM L-Glutamine (Invitrogen), 50 units/ml Penicillin and 50 ug/ml Streptomycin (Invitrogen). Immediately after removal thymi were incubated for 15 min at room temperature in 2 mg/ml papain (Worthington Biochemical, Lakewood NJ) in RPMI-1640. The Papain reaction was stopped by adding 1 volume of TEC-10 media and the tissue was then broken up by trituration. Cells were then plated on a 10 cm tissue culture plate (Nunc) in 10 ml of TEC-10. The cells were allowed to grow for 3 days in original media. After three days the media was changed to eliminate thymocytes. Cells were maintained in culture for fewer than 7 days and were not passaged prior to Nucleofection.

For primary culture of postnatal TECs, thymuses were removed from 3 day old or 30 day old Swiss Webster mice. After removal, thymuses were placed in TEC-10 medium and allowed to settle in a 15 ml tube for 5-10 mins. The thymuses were rinsed in three successive changes of TEC-10 medium to remove non-thymic cell types clinging to the surface. Cells were isolated by digesting the thymus tissue with 0.2 mg/ml collagenase D in RPMI-1610 containing 2% fetal bovine serum, 20mM HEPES at 25°C for 20 mins. The tissues was then incubated in a medium containing RPMI-1610, 2% fetal bovine serum, 20mM HEPES, 0.2 mg/ml collagenase, 0.2 mg/ml dispase and 25μg/ml DNase I for 40 mins at 37°C. Five volumes of TEC-10 medium was added to stop the enzymes and the tissue clumps were broken up by trituration. The resulting cell suspension (from 8–11 thymuses) was plated onto a 10 cm tissue culture plate.

### Nucleofection of TEC lines and primary thymic epithelial cells

Nucleofection of the thymic epithelial cell lines and primary TECs was performed using the Amaxa Nucleofector II device. Based on technical advice from Amaxa Inc. support staff we used the cell line Nucleofection kit V to optimize conditions for the 4 TEC lines. Seven samples each containing 2 × 10^6^ cells were prepared for each cell line and each sample was Nucleofected using one of 6 different programs (A-20, T-20, T-30, X-01, X-05, D-23) with the untransfected sample serving as a control. Nucleofections were performed according to Amaxa’s recommendations with 5μg of pmaxGFP. The pmaxGFP plasmid contains a CMV promoter driving a Pontellina (copepod) green fluorescent protein coding sequence [58] and an SV40 poly A site (Amaxa News #3). After each sample was transfected it was immediately transferred to 1 ml of culture media that had been warmed to 37°C and placed in an incubator (37°C, 5% CO_2_) until all samples were ready for plating. For each sample, half of the cells were plated and the medium was changed after 3-5 h to remove dead cells. The plates were incubated for 24 h and then fixed in 1% paraformaldehyde in 1X PBS and stored at 8° C for flow cytometry analysis. The other half of each sample was stained with propidium iodide (Sigma) immediately after Nucleofection for flow cytometry analysis of cell mortality due to the procedure.

Primary thymic epithelial cells were maintained in TEC-10 media in 10 cm tissue culture dishes (Nunc). For optimization experiments the primary TECs were divided into 5 samples of 2 × 10^6^ cells each. The samples were suspended in 100μl of solution V (Amaxa Inc.) containing 1μg of pmaxGFP plasmid and subjected to one of four Nucleofector programs as recommended for primary epithelial cells (S-05, T-13, T-20, X-05). One sample was not transfected and served as a negative control. Each sample was seeded into one well of a 6 well tissue culture plate (Nunc) and allowed to grow for 24 hours before observing GFP expression. For cell counts, the transfected cells were plated on microscope slides in 250 μl droplets and allowed to attach for 24 hours.

#### Cell counting and flow cytometry

The efficiency of pmaxGFP transfection of the cell lines was measured using a FACScalibur cytometer (Becton Dickenson) and the data were analyzed using FloJo software. Cell death resulting from each transfection was measured by counting the number of cells that were stained by propidium iodide in each sample immediately after Nucleofection. Propidium iodide stock solution (1 mg/ml Sigma) was diluted in 1X PBS to 1μg/ml. Medium was removed from each sample by centrifugation. The cell samples were resuspended in 500μl of propidium iodide staining solution (1μg/ml propidium iodide in PBS) and allowed to incubate at room temperature for 30 minutes. The cells were pelleted, resuspended in cold PBS and analyzed by flow cytometry within 30 mins.

The transfection efficiency of the primary cells was measured by counting cells in multiple microscope fields of transfected samples and untransfected control samples. After 24 hours of culture, half of the samples were fixed with acetone and cytokeratins were detected using a pan-cytokeratin polyclonal primary antibody (Dako) and a secondary antibody conjugated to texas red (Jackson ImmunoResearch). The remaining samples were not fixed with acetone so that GFP expression could be observed for counting. All samples were stained with DAPI using DAPI containing aqueous mounting media (Invitrogen) so that nuclei could be visualized for cell counts. To measure GFP expression 10 micrographs were taken of each sample and the cells were counted. To measure viability, mean total cell numbers in transfected samples and in untransfected controls were determined.

## Authors’ Contributions

R.T.O. and Q.W. performed the experiments. R.T.O. and B.G.C. wrote the manuscript. B.G.C. conceived and designed the study.

## Acknowledgements

We thank Ms. Liz Horton at Amaxa Inc for technical advice and Dr. Nancy Manley for reading the manuscript. We also thank Dr. Shinyun Xiao for assistance with flow cytometry. This work was supported by startup funds (to B.G.C.) from the University of Georgia.

